# Opto-MASS: a high-throughput engineering platform for genetically encoded fluorescent sensors enabling all-optical *in vivo* detection of monoamines and opioids

**DOI:** 10.1101/2022.06.01.494241

**Authors:** Michael Rappleye, Adam Gordon-Fennel, Daniel C. Castro, Avi K. Matarasso, Catalina A. Zamorano, Carrie Stine, Sarah J. Wait, Justin D. Lee, Jamison C. Siebart, Azra Suko, Netta Smith, Jeanot Muster, Kenneth A Matreyek, Douglas M. Fowler, Garrett D. Stuber, Michael R. Bruchas, Andre Berndt

## Abstract

Fluorescent sensor proteins are instrumental for detecting biological signals *in vivo* with high temporal accuracy and cell-type specificity. However, engineering sensors with physiological ligand sensitivity and selectivity is difficult because they need to be optimized through individual mutagenesis *in vitro* to assess their performance. The vast mutational landscape proteins constitute an obstacle that slows down sensor development. This is particularly true for sensors that require mammalian host systems to be screened. Here, we developed a novel high-throughput engineering platform that functionally tests thousands of variants simultaneously in mammalian cells and thus allows the screening of large variant numbers. We showcase the capabilities of our platform, called Optogenetic Microwell Array Screening System (Opto-MASS), by engineering novel monoamine and neuropeptide *in vivo* capable sensors with distinct physiological roles at high-throughput.

## Introduction

Genetically encoded fluorescent indicators (GEFIs) are protein-based sensors that increase fluorescence intensity upon target ligand binding[1]. The basic engineering principle combines a ligand-specific binding domain with a fluorescent reporter protein and tunes their connection by mutating the amino acids linking the two domains. Recently, a new subset of GEFIs was constructed by grafting a circularly permuted fluorophore (cpGFP) into the third intracellular loop of dopamine G-protein Coupled Receptors (GPCRs) to engineer dopamine sensors (DA) [2], [3]. As with most GEFIs, several hundred sensor variants were screened to optimize signal amplitudes and dopamine detection. However, the screened mutations represent a small fraction of the 160,000 variants that constitute the mutational landscape of the four targeted residues (20^4^). Better sensors can likely be identified if screening methods could screen more variants. The grafting principle has been demonstrated to work on a host of GPCRs, expanding the available sensors to include acetylcholine, serotonin, norepinephrine, and orexin [4]–[7]. But similarly, protein engineering bottlenecks tested only a few hundred variants during optimizing ligand sensitivity and signal output.

Traditionally, engineering fluorescent biosensors require a multistep, resource-intensive process. Researchers generate individual mutations in plasmid DNA by PCR, purify them from *E*.*coli* one by one, and express the variants in a heterologous expression system for analysis. Membrane-bound GPCR-based sensors require testing in mammalian host cells such as HEK293 cultures because yeast and bacteria cells have difficulty expressing a diversity of fully functional GPCRs at their membranes[8]. On the other hand, to test constructs in mammalian cell cultures, researchers transfect individual plasmids into cells seeded in multi-well plates (24-384 wells), limiting throughput. The fluorescent output of sensor variants is then tested upon ligand application, often under saturating conditions to elicit maximum responses. The mutation and screening process must be repeated hundreds of times, as performance is notoriously difficult to predict in these highly dynamic fusion proteins. The field needs to address the significant gap presented by resource-intensive techniques currently used to engineer sensors for the wealth of GPCRs in mammalian physiology because current methods preclude the development of sensors for most GPCRs.

Here, to address this gap, we present a high throughput platform to construct genetically encoded fluorescent indicators rapidly. Furthermore, the mammalian host cells are engineered to express one single variant while using commercial transfection reagents. We physically separate the individual sensor expressing cells into single wells in a microwell array. Taken together, we functionally screen hundreds of cells simultaneously, resulting in thousands of tested cells per day. We quickly rank the sensor’s phenotypes in real-time using fluorescence microscopy and automated image analysis. We can then faithfully identify the best-performing variants from thousands of cells within minutes. When considering the engineering of GEFI optimization platforms, we must design the platform for broad application, ease of adoption, and throughput. Protein engineering pipelines using mammalian cells have been applied to optimize genetically encoded voltage and calcium indicators [9], [10]. However, the functional screening of sensors was either limited in throughput or did not provide signal readouts under dynamic conditions. To showcase the broad applicability of our pipeline, we engineered *in vivo* capable monoamine and opioid sensors. In summary, the OPTO-Mass platform addresses a gap in the field by providing a higher throughput platform to optimize GPCR-based biosensors to *in vivo* capabilities at a faster throughput and reduced resource commitments.

## Results

### Opto-MASS Design goals

We envisioned a method that functionally screens thousands of optogenetic sensor variants each day to increase engineering throughput significantly. New, improved sensors could then be used as a scaffold for iterative mutations and further optimizations using the platform (Fig. 1). We identified four necessary features to achieve this goal. 1.) A single-step library generation strategy to make an extensive, unbiased library of sensor variants in DNA plasmids. 2.) A mammalian expression system wherein one single plasmid is expressed per cell while maintaining high transfection efficiency. 3.) The ability to read out functional, dynamic signals from hundreds of cells simultaneously under physiological conditions. 4.) Recovery of the genetic content that encodes high-performing variants.

**Figure 1:**
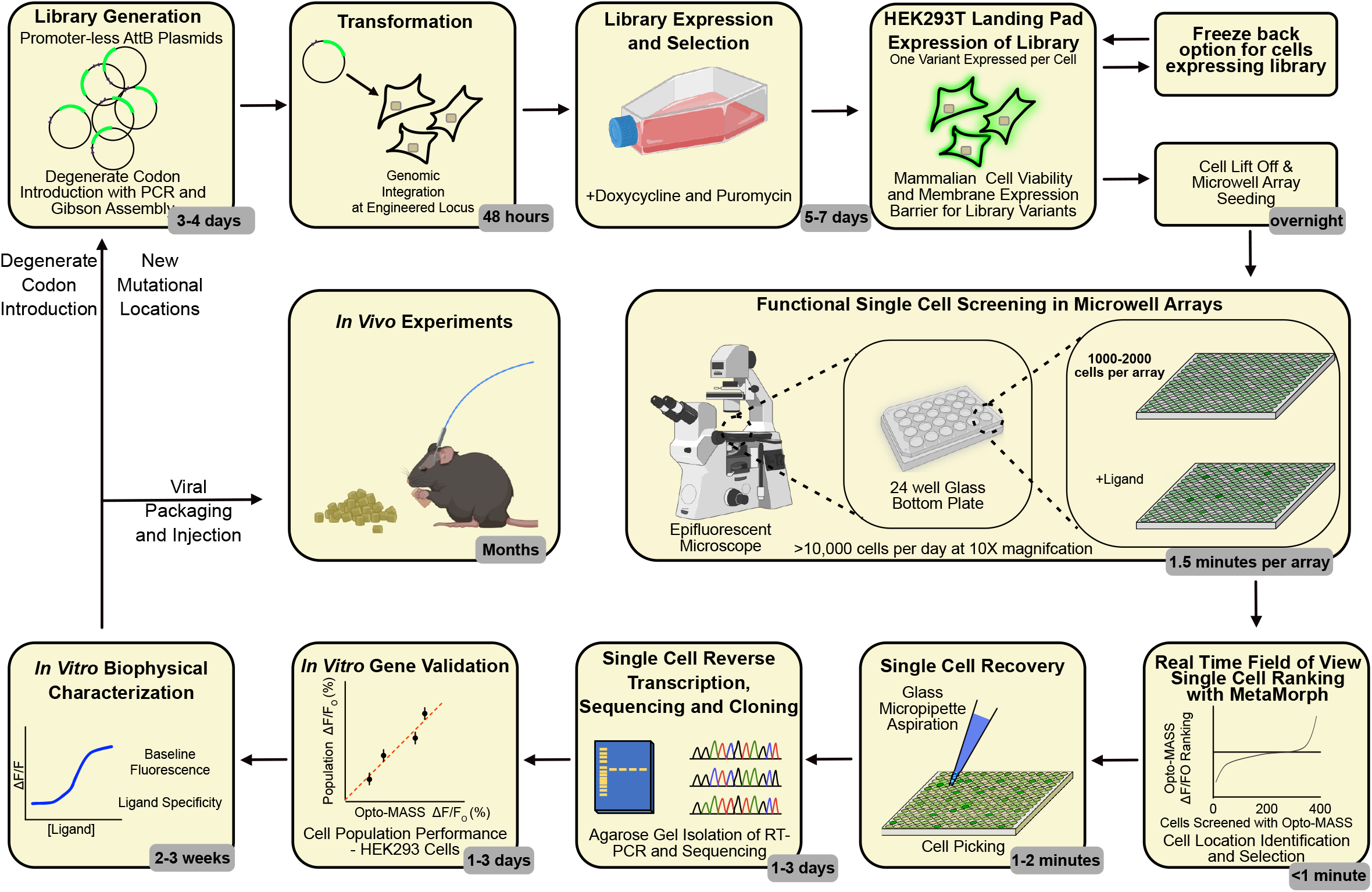
The Optogenetic Microwell Array Screening System (Opto-MASS) Researchers need a facile, rapid, and scalable system to build optogenetic sensors. Opto-MASS combines microwell array screening technology and mammalian genetic expression systems to rapidly screen thousands of cells/variants of sensors on a platform that can complete sensor engineering in less than a month. First, a randomized mutational library of the sensor is constructed in landing pad compatible plasmids. The library is then recombined in the HEK293T “landing pad” cell genome into a single locus. After doxycycline induction and puromycin selection for 5-7 days, the library can be frozen back for later screening or seeded onto microwell arrays. The Opto-MASS microwell arrays are screened on an inverted fluorescent microscope, and sensors are ranked nearly instantaneously in each field of view. After identifying the highest-performing cells, a glass micropipette is used to aspirate them physically. Next, we use RT-PCR to identify the underlying sensor gene. The recovered gene is then transfected into HEK293WT cell cultures to characterize the sensor’s biophysical phenotype in detail on a population level. If the sensor retains the desired characteristics, we can package it into viral vectors for *in vivo* experiments or use the sensor as a template for a new round of library screening.

To achieve these goals, we put the ‘landing pad’ HEK 293T TetBxb1BFP cells at the center of our platform (a kind gift from Dr. Douglas Fowler) [11]. They enable the facile expression of a single variant per cell in mammalian cells. The sensor expressing cells are screened in customized PDMS (polydimethylsiloxane) microwell arrays placed in 24-well cell culture plates (Fig. 1). The microwell arrays physically separate the cells to enable an easy, functional readout of fluorescent signals from hundreds of cells simultaneously. The cells on each array are ranked based on ligand-dependent fluorescence changes in real-time. We physically recover the cells by aspirating them with a glass micropipette controlled by a micromanipulator. Next, we perform single-cell RT-PCR on the recovered cell to identify the sensor encoding gene. We validate the recovered gene’s phenotype by biophysical characterization in cultured HEK293 cell populations. After assessing the sensor’s performance, we could iterate the process and use the recovered variant as a scaffold for the next library or package the sensor into a virus for *in vivo* experiments.

### Landing pad mammalian expression system

The landing pad expression system has several key features that make it an amenable solution to our design goals. The plasmids encoding for sensor variants contain no constitutive mammalian promoter. Instead, upstream of our genes of interest, an AttB recombination site enables irreversible recombination with an AttP site inserted into the genome. We cloned a tricistronic gene downstream of the AttB recombination site on our sensor library plasmids. The three genes encode the green fluorescent GPCR sensor, red fluorescent mCherry (control for image analysis), and a puromycin resistance gene (selection of recombined cells) and are each separated by self-cleaving P2A sequences (Fig. 2A). Thus, after Bxb1 mediated recombination into the engineered locus, each protein functions independently. A Tet inducible promoter drives the expression of the tricistronic cassette, allowing the tuning of genetic expression (4-10 μg/mL doxycycline). As a result, while transfection reagents can introduce more than one plasmid per cell, only one plasmid can recombine into the genomic landing pad. Thus, only a single variant is expressed per cell (Fig. 2B).

**Figure 2:**
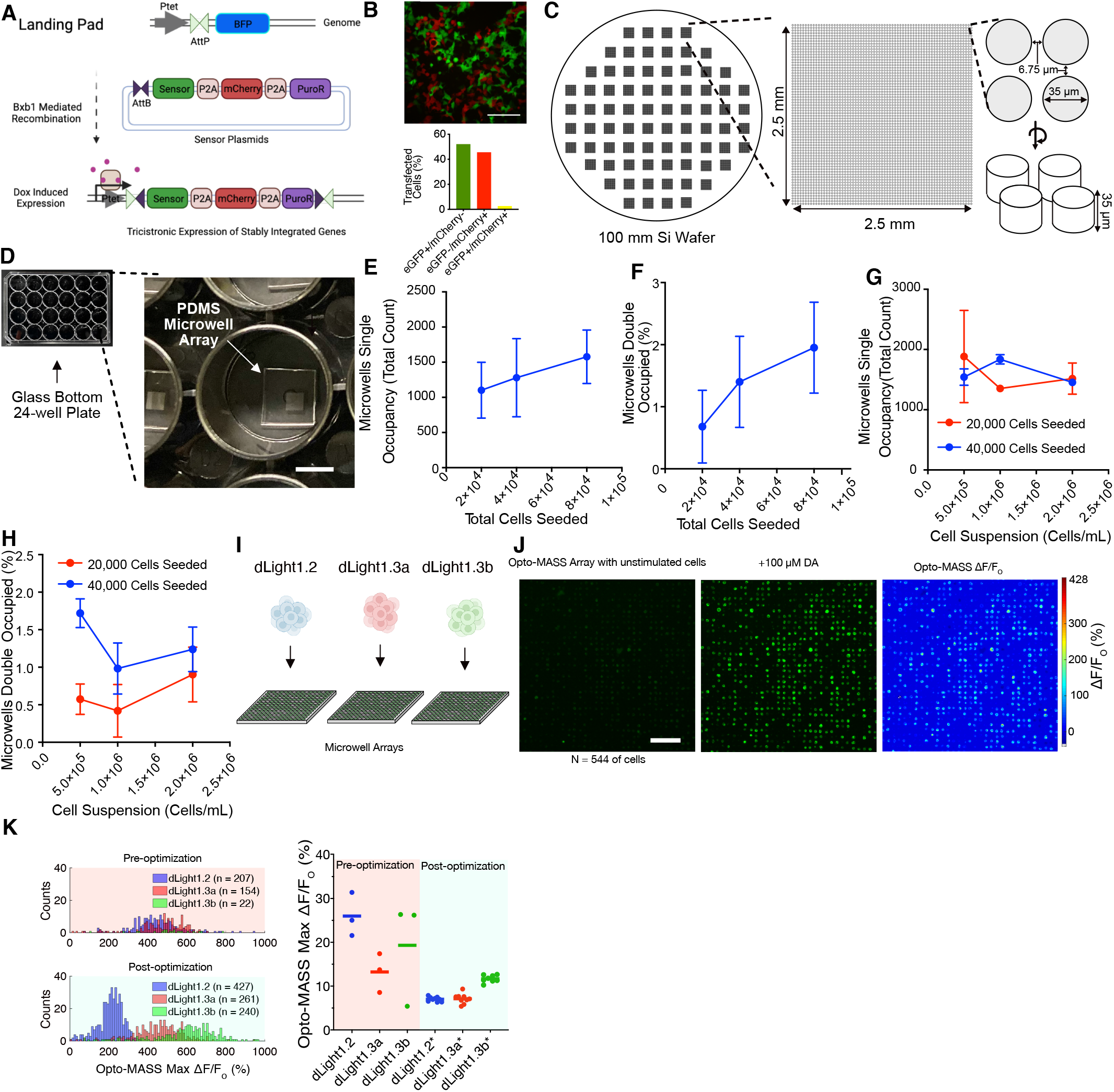
Design and Engineering of an Optogenetic Microwell Array Screening System. A) Schematic of the genomic landing pad in the HEK293T landing pad cells (HEK 293T TetBxb1BFP) pre and post-sensor plasmid integration. B) Representative false-color image of HEK293T TetBxb1BFP cells transfected with a mix of mCherry and eGFP plasmids (upper panel, 100 μm scale bar). Summary data for the field of view for fluorophore expression (lower panel). The majority of cells express only one fluorophore. C) Silicon master mold of the microwell arrays (left), dimensions of one array (middle), and individual micro-arrays (right). The Si wafer was etched using Deep Reactive Ion Etching. D) Polydimethylsiloxane (PDMS) microwell arrays are cut from the master mold after curing, and BSA nanostamping and are placed ‘wells up’ in the bottom of a 24-well glass bottom plate for cell seeding and screening. E-H) HEK293T landing pad cells expressing dopamine sensors are seeded onto separate microwell arrays at varying total amounts to determine the optimal seeding density. The total amount of cells seeded was varied to determine if it increased well occupancy (E and F), or cell suspension concentration was altered to determine effects on well occupancy rates (G and H). I) For initial testing, we seeded three distinct dopamine sensors with know signal amplitudes onto separate arrays. J) Representative stimulation of an isogenic population stimulation of landing pad cells expressing dLight1.3b and stimulated with 100 μM dopamine on the Opto-MASS array. N = 544 cells in the field of view. 200 μm scale bar. K) Array stimulation optimization. Cell seeding density, cell handling, microscope focusing, and array fabrication were optimized to reduce the coefficient of variation in arrays stimulation of three different dopamine sensors. Saturating concentrations of dopamine were used. Average CoV pre-optimization 19.52% (n = 9 microwell arrays tested) were lowered to 8.49% (n = 26 microwell arrays tested).

### Microwell array design and cell seeding

The design goal for the microwell array was to image thousands of physically separated cells within a single field of view, straightforward analysis of sensor function, and physical recovery of sensor variants. The microwell arrays were designed to accommodate a single HEK293T cell. The physical separation provided by the microwells expedites the automated analysis of fluorescent signals from individual cells (Fig. 1). We fabricated a silicon master mold from a 100mm silicon wafer (University Wafer, Fig. 2C) using deep reactive ion etching (DRIE) to etch away the wafer and generate a negative mold of the arrays. We varied well diameter (20-100 µm), distance (5-10 µm), and depth (20-50 µm) for initial prototyping. Our goal was to reach maximum well density in the field of view of our camera (Photonics Prime 95B 2048×2048 pixels at 11 µm per pixel) while holding only one cell per well. The array size is 2.5 × 2.5 mm to match the field of view of our camera at 5X magnification. We found optimal well parameters at 35 µm diameter, 6.75 µm distance, and 35 µm depth resulting in 3600 wells per array. This enables the observation of a maximum of 3600 wells at 5X magnification or 900 wells at 10X. Each 100mm silicon wafer carries 76 microarrays (Fig. 2C). We use polydimethylsiloxane (PDMS, Sylgard 184) to cast our Opto-MASS arrays by standard soft lithography techniques. (Fig. 2D). We chose PDMS due to its optical clarity, low cost, and low cytotoxicity on our experimental timescales (12-16 hours). After curing the PDMS on the master mold, the PDMS slab was nanostamped with BSA to reduce cell adhesion outside of the microwells during initial cell seeding (see methods).

After removing dust and debris from the back of the PDMS slab, individual arrays were cut from the PDMS slab and placed in the wells of a 24-well, glass-bottom dish (CellVis, NC0397150, Fisher Scientific). We optimized cell suspension seeding conditions for the HEK293T landing pad cells to maximize well occupancy while reducing the occurrence of multiple cells in a microwell (Fig. 2E-H). For our library screening experiments, we chose to seed 80,000 HEK 293T cells at a concentration of 0.5×10^6^ cells/mL per array. This results in single occupied wells at about 30%-50% total occupancy of arrays routinely providing up to 1800 observable cells at 5X or 300-500 cells at 10X magnification.

### Automated Image Acquisition and Analysis Protocol

During the functional screening of the libraries, we required an efficient way to track and rank the cell responses upon adding ligands. Our microscope and imaging setup was operated using MetaMorph imaging software (Molecular Devices). The control fluorophore, mCherry, was imaged before and after stimulation to remove any cells that moved into or out of the field of view (Supp.Fig. 1A-C). During stimulation, the cells were imaged continually under GFP wavelengths (EX: 474/27 nm, EM: 520/35 nm), and ligands were added to the bath to screen for sensor functionality (Supp. Fig. 1D). We used mCherry to define regions of interest (ROIs) for automated analysis in MetaMorph. ROIs were then measured for size and excluded if they were too large or too small for typical cells. (Supp. Fig. 1 and 2).

The ROIs were transferred to the pre-stimulation mCherry image to measure mean grayscale values as a control for fluorophore expression. The ROIs were then transferred to an image stack covering the stimulation period. The ROI’s average GFP grayscale value (i.e. fluorescence intensity) was measured and exported to Excel for offline analysis (Supp. Fig. 1D and 1E). Next, the stimulation image stack was split into two different stacks, the pre- and post-stimulation stacks, with the average fluorescence intensity projection taken for both stacks. The resulting images were then divided and multiplied by 1000, so that ROIs that increased in fluorescence had a value higher than 1000, and those that decreased had a value lower than 1000 (Supp. Fig 1F). We dubbed the ratio value the Opto-MASS Ranked Ratio for each ROI and it was used to identify ROIs that had the greatest increase in fluorescence in the field of view (Supp. Fig. 1F).

We excluded images close to the ligand application because they could cause motion artifacts that alter the fluorescence calculation (Supp. Fig. 2D). After identifying the ten highest-ranked ROIs, we recovered the corresponding cells under bright-field. (See Supp. Fig. 2 for detailed task execution during screening).

### Opto-MASS screens functionally diverse sensor populations

Next, we demonstrated that the platform could correctly identify signals from sensors with different but known signal amplitudes under similar ligand concentrations. For this purpose, we cloned the dopamine (DA) sensors dLight1.2, dLight1.3a, and dLight1.3b into landing pad plasmids[3]. We generated isogenic populations of the sensors by stably integrating the different dopamine sensors into separate landing pad cell populations, inducing their expression with doxycycline (10 μg/mL) and selecting for them with puromycin (0.75-1μg/mL) (Fig. 2I). We then seeded the isogenic populations onto different arrays (i.e. each cell population expressing one sensor on separate arrays) to test for sensor functionality (Fig. 2I-K).

We expected that the dopamine-dependent fluorescent changes at saturating DA concentrations (100 μM) should create differentiable responses between these sensors. We achieved this goal after extensive protocol optimization, including expression time, recovery on PDMS arrays, cell seeding density, and PDMS thickness. Using the optimized protocol, we could distinguish the different dopamine sensor populations based on their average fluorescent output (Fig. 2K). Importantly, we yielded a reduced coefficient-of-variance for the sensor populations on the arrays to demonstrate the signal readout was representative of the underlying sensor (Fig. 2K). The significantly improved CoV of approximately 10% ΔF/F_o_ provides a reduced window for potential outliers and cell-to-cell variability (Fig. 2K)[9].

### Using Opto-MASS’s enhanced screening capabilities to identify a high-performance monoamine sensor

Next, we demonstrated that we could identify variants with optimized signal amplitude and ligand sensitivity variants from a large mutational library of an existing sensor framework. We chose the dopamine sensor dLight1.1 for this purpose because of its reliable signal generation and utility in neuroscience [3]. We targeted the four residues flanking either side of the fluorophore cpGFP inserted into the human D1 dopamine receptor (Fig. 3A). These four residues have been demonstrated to play a critical role in coupling fluorophore brightness to ligand-dependent changes in the receptor domain, but only a small fraction of the possible mutational space (20^4^) has been investigated in the literature so far [3], [12]. We built a library of randomized mutations at these sites by incorporating DNA primers with degenerate codons (IDT) in a single PCR step and subsequent Gibson Assembly (NEB) (Fig. 3A, Supp. Fig. 3, > 200,000 *E*.*coli* transformants). Sampling DNA sequences from 23 *E*.*coli* colonies revealed a relatively even distribution of four nucleic acids at the targeted sites (Fig. 3B). We transfected the randomized plasmid library into a population of 250K cells. The landing pad cells had been transfected with plasmids encoding for a nuclear-localized Bxb1 recombinase twenty-four hours prior (Fugene6, Promega). We drove library expression and selection by adding doxycycline (10 μg/mL) and puromycin (1 μg/mL), respectively, after 24h. After two days of selection, cells were combined and passaged into a T25 flask. Cells cultures were expanded for 5-7 days until the selection was complete. Next, we lifted the cells with Trypsin (0.05%) and EDTA from the flasks and seeded cells onto microarrays placed in glass-bottom 24-well plates. We let the cells recover in incubators for 12-16 hours overnight before screening (37°C, 5% CO_2_). Plates were imaged under epifluorescence using the GFP channel (474 nm excitation, 520 nm emission, 500 ms exposure) for dopamine signals and the mCherry (578 and 641 nm) channel as negative controls. In one trial, we screened ∼13,000 cells (Fig. 3C). Cells on each array were stimulated by a low 500 nM dopamine application via an automated syringe pump at consistent time points. The simultaneous increase of fluorescent signals from the cells demonstrates that diffusion of the dopamine was immediate (Supp. Fig. 2D). We could faithfully rank cells in each field of view in real-time using customized MetaMorph scripts (Supp. Fig 2&3). We recovered the highest-ranking cells from the arrays using glass micropipettes connected to a syringe for aspiration and controlled by a micromanipulator. Each cell was placed into separate microcentrifuge tubes containing Tris/EDTA buffer and the reducing agent dithiothreitol (DTT 2.44 mM) to prevent RNA degradation by RNAse.

**Figure 3:**
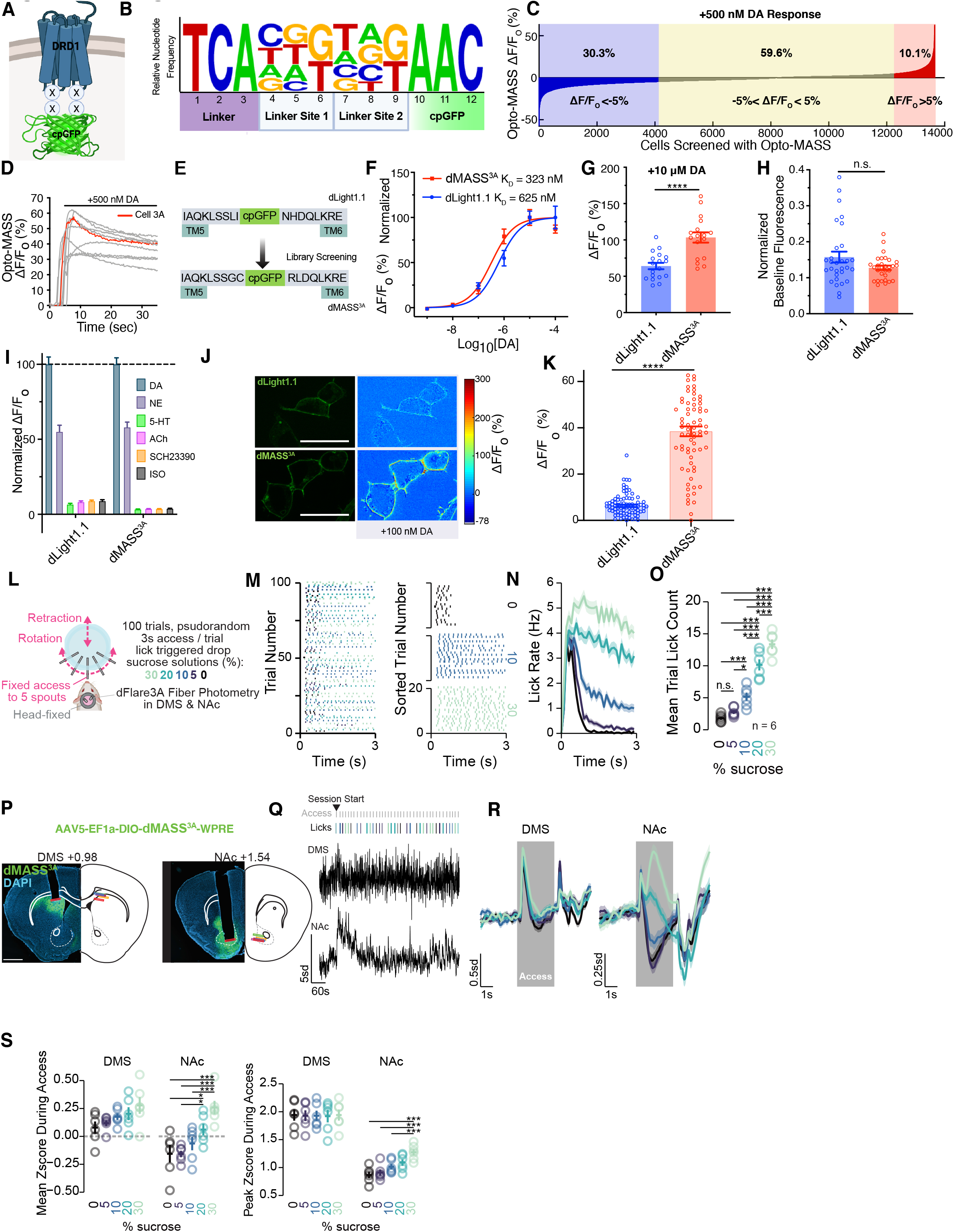
Opto-MASS enhances an *in vivo* capable monoamine sensor. A) Schematic of the residues targeted for generating a mutational dopamine sensor library from dLight1.1 (denoted by X). The four residues are located in the linker between the cpGFP and GPCR DRD1 domain and were targeted by site-saturated randomized mutagenesis. B) Summary of 23 sampled sequencing results from the c-terminal linker reveals near equal distribution of all possible nucleotides at the targeted sites. C) Aggregate responses from one screening session of the dopamine sensor library. 13,656 cells were screened in one day. 30.3% of the screened cells had a dopamine (DA) dependent decrease in fluorescence (ΔF/F_o_ < -5%) 59.6% had no change in fluorescence and 10.1% had an increase in fluorescence (ΔF/F_o_ > 5%). D) Exemplary responses of selected cells from the screen. Cell 3A, isolated to become variant dMASS^3A^, is highlighted in red. E) dMASS^3A^ mutations in both linker regions were identified by RT-PCR and Sanger sequencing post-screening and compared to dLight1.1. F) Comparison of the apparent dopamine affinity curves of the parent scaffold dLight1.1 and dMASS^3A^ under epifluorescence microscopy. (Single site binding n = 3 wells, 6 cells/well). G) dLight1.1 and dMASS^3A^ response to 10 μM DA in HEK293 mammalian cell cultures. (p < 0.0001, unpaired t-test, n = 5-6 cells/well, 3 wells). H) Baseline fluorescence of dLight1.1 and dMASS^3A^ normalized to C-terminal tagged mRuby3 (negative control). (p = 0.0734, unpaired Student’s t-test). I) Pharmacological specificity of dLight1.1 and dMASS^3A^ to 10 μΜ of compounds, normalized to 10 μM DA stimulation(dLight1.1 100 ± 4.85%, dMASS^3A^ 100±4.29), NE is norepinephrine(dLight1.1 55.06 ±4.31 dMASS^3A^ 58.00±3.46%), 5-HT is 5-hydroxytryptamine (dLight1.1 6.61± 0.69% dMASS^3A^ 3.32± 0.20%), ACh is acetylcholine (dLight1.1 8.22±0.58%, dMASS^3A^ 3.61 ±0.14%), SCH23390 (D_1_ receptor antagonist dLight1.1 8.87±0.58%, dMASS^3A^ 3.58±0.16%), Iso is isoproterenol (dLight1.1 8.89±0.83%, dMASS^3A^ 3.80±0.21%). N = 40 cells. All plots mean ± SEM. J) Representative confocal images of HEK293 cells expressing dopamine sensors and membrane-localized responses to 100 nM dopamine bath application. 30 μm scale bar. K) Average dLight1.1 and dMASS^3A^ membrane-localized responses in HEK293 cells to 100 nM dopamine imaged on a confocal microscope. L) Training RPE paradigm for pseudorandom access to sucrose solutions of varying concentrations. Over 3 sessions, mice were given 100 trials of 3s access to 5 concentrations of sucrose (colors in subsequent plots indicate the concentration of sucrose). M) Consummatory licks during 3s access to each concentration of sucrose displayed as mean binned licking over time (N) and mean total lick count during access for each concentration (O). P) Histological images of dMASS^3A^ expression and fiber tip placement in the dorsal medial striatum and Nucleus Accumbens. 1 mm scale bar. Q) Representative fiber photometry trace of dMASS^3A^ fluorescence simultaneously recorded in the DMS and NAc. Grey lines indicate 3s access periods, and lick lines indicate licks for each concentration. R) Mean fluorescence traces in the DMS (left) and NAc (right) over time during access to each concentration (ribbon depicts SEM). S) Mean (left) and peak (right) fluorescence during the access period shows clear scaling in the NAc but not in the DMS.

Using a gene-specific primer, we recovered the sensor encoding gene from each cell by RT-PCR (SSIV First-Strand Synthesis, ThermoFisher). PCR amplified cDNA was recovered from agarose gels following electrophoresis and identified by routine Sanger Sequencing (GeneWiz). The recovered sensor sequences were cloned into mammalian expression vectors (pC_DNA3.1, CMV promoter) for subsequent transfection (Lipofectamine 3000, ThermoFisher) and biophysical characterization into HEK293WT cell cultures.

Importantly, the high-throughput capabilities of the platform allowed us to identify highly dynamic variants at lower ligand concentrations instead of being biased towards maximum brightness under saturating conditions. During screening, we observed approximately one third of our measured cells decrease in brightness upon ligand addition (30.3% ΔF/F_o_ < -5%), 59.6% had no fluorescence response (−5%< ΔF/F_o_ < 5%), and 10.1% of the population increased in fluorescence (ΔF/F_o_ > 5%).

We observed several high-performing variants (Fig. 3D) and recovered cell 3A to pursue further characterization in HEK293 cell populations. We dubbed the recovered variant dMASS^3A^. All the targeted linker sites were mutated compared to dLight1.1 in dMASS^3A^ (Fig. 3E). When screened with epifluorescent microscopy, dMASS^3A^ had a lower K_D_ than the parent construct, dLight1.1 (K_D_ = 323 nM and 625 nM, respectively, Fig. 3F). dMASS^3A^ had 1.6-fold greater response than dLight1.1 at 10 μM DA (p< 0.0001, Student’s T-test, unpaired, Fig. 3G). The baseline fluorescence of dMASS^3A^ was close to the parent construct, dLight1.1 (p = 0.0734, unpaired Student’s t-test, Fig 3H).

The increased fluorescence output came at no apparent loss of molecular specificity of the sensor (Fig. 3I). To align our measurements with previous studies, we tested dMASS^3A^ on a confocal microscope at low dopamine concentrations (Fig. 3J-K >6 fold, p<0.0001, two-tailed t-test). Here, dMASS^3A^ also outperforms the parent construct, dLight1.1, demonstrating sensors can be enhanced in specific ways by targeting Opto-MASS screening conditions towards the desired sensor characteristics.

### dMASS^3A^ detects Dopamine signals *in vivo*

Due to the broad utility of GPCR-based sensors in neuroscience, we aimed to demonstrate that sensors engineered by our pipeline are compatible with neuronal *in vivo* and *in vitro* detection methods. Here, we validated dMASS^3A^ *in-vivo* within the dorsal medial striatum (DMS) and nucleus accumbens (NAc) of mice during consummatory behavior using fiber photometry (Fig. 3L). Dopamine signaling in the DMS and NAc are essential modulators of operant and consummatory behavior[13], [14]. Still, the precise dopamine signaling dynamics in these structures during free consumption of different reward magnitudes remain unclear. To investigate this, we recorded dopamine release using dMASS^3A^ using fiber photometry during limited windows of free-access consumption of 5 concentrations of sucrose (Fig. 3L). We trained head-fixed mice to lick and consume sucrose during 100 trials of 3s access (Fig. 3K-N). Mice exhibited consumption of sucrose that was dependent on the concentration of sucrose which was manifested as more licking for high concentrations of sucrose compared to low concentrations (Fig. 3N-O; F_4,20_ = 114.63, P = 1.72e-13). dMASS^3A^ was strongly expressed in the NAc and DMS (Fig. 3J) and showed explicit dynamics during behavior (Fig. 3O-R). In the DMS, dMASS^3A^ signals showed strong, transient increases in response to the onset of the access period that did not scale with the concentration of sucrose. The transient increase in dMASS^3A^ fluorescence in the DMS coincides with the onset of licking behavior, implying that dopamine release in the DMS may initiate but not sustain motor actions of consumption [15], [16]. On the contrary, in the NAc, dMASS^3A^ signals showed a two-component response with an initial rise at the onset of the access period and a secondary rise or drop in signal during the middle of the access period (Fig. 3P). The mean and peak dMASS^3A^ fluorescence in the NAc showed explicit scaling with the concentration of sucrose with higher fluorescence during consumption of higher concentrations of sucrose and decreases in fluorescence during access periods with lower concentrations of sucrose. The change in dMASS^3A^ fluorescence in the NAc does not directly track licking behavior, as the dMASS^3A^ response begins to return to baseline before licking returns to 0 (Fig. 3O and P). Instead, the dMASS^3A^ response in the NAc may represent an initial detection response and a secondary value signal related to sucrose concentration during the access period[17].

### Opto-MASS engineers a neuropeptide sensor capable of *in vivo* detection of opioids

For the final validation of the capabilities of our pipeline, we chose to engineer a GPCR-based sensor framework that currently lacks *in vivo* detection capabilities. We selected to optimize a sensor prototype called mLight based on the Mu-opioid GPCR (MOR)[3]. Neuropeptides such as endogenous opioids are hypothesized to function through low concentration volume transmission instead of fast, high concentration synaptic transmission like monoamine neurotransmitters[18]. The low concentrations make *in vivo* neuropeptide detection more difficult. Endogenous opioid peptides include endorphins, enkephalins, dynorphins, and nociceptin, which help regulate motivation, stress, reward, gastrointestinal mobility, hedonic homeostasis, feeding, and other behaviors through the opioid receptors [19]–[22]. The opioid peptides bind to different subtypes of opioid receptors (mu, delta, kappa, nociceptin) with varying sensitivity but are rarely exclusive to just one target receptor [23]. Current techniques to monitor opioid peptide release either lack cell-type specificity, the kinetics to link signaling events with animal behavior, or are incompatible with current *in vivo* imaging technologies[24]–[26]. Previously, researchers have inserted the cpGFP moiety into the ICL3 of the mu-opioid receptor (MOR) to make a prototype opioid GEFI mLight. However, it suffered from poor dynamic range and cell surface expression, precluding it from *in vivo* use [3].

First, we validated that the published cpGFP domain insertion location in mLight was the most optimal position (Supp. Fig. 4A-B). We then enhanced membrane trafficking in the landing pad cell system by adding membrane trafficking and ER export sequences (TS-ER) to the sensor scaffold (Supp. Fig. 4C)[27]. Next, to increase the allosteric coupling between the two domains, we targeted mutations to four residues within the linkers between MOR and cpGFP (Fig. 4A, Supp. Fig. 4D). We screened >23,000 cells/variants at 1 μM [D-Ala^2^, N-MePhe^4^, Gly-ol]-enkephalin (DAMGO), a synthetic enkephalin with high specificity for the MOR. We recovered a high-performing variant from the library for testing in HEK293 cell populations and dubbed it μMASS^2A^. μMASS^2A^ had significantly better responses to 500 nM (∼4.6 fold) and saturating concentrations of DAMGO (∼3.8 fold) than the parent construct mLight (Fig. 4C and D). Similar to the native MOR, the sensor could detect several types of opioid peptides, such as the Met- and Leu-enkephalins, dynorphin A and beta-endorphin with differing apparent affinities (Fig. 4E, DAMGO K_D_ = 243 nM, Methionine-enkephalin K_D_ = 99 nM, leucine-enkephalin K_D_ = 637 nM, β-endorphin K_D_ = 1176 nM, Dynorphin A K_D_ = 1125 nM [23]).

**Figure 4:**
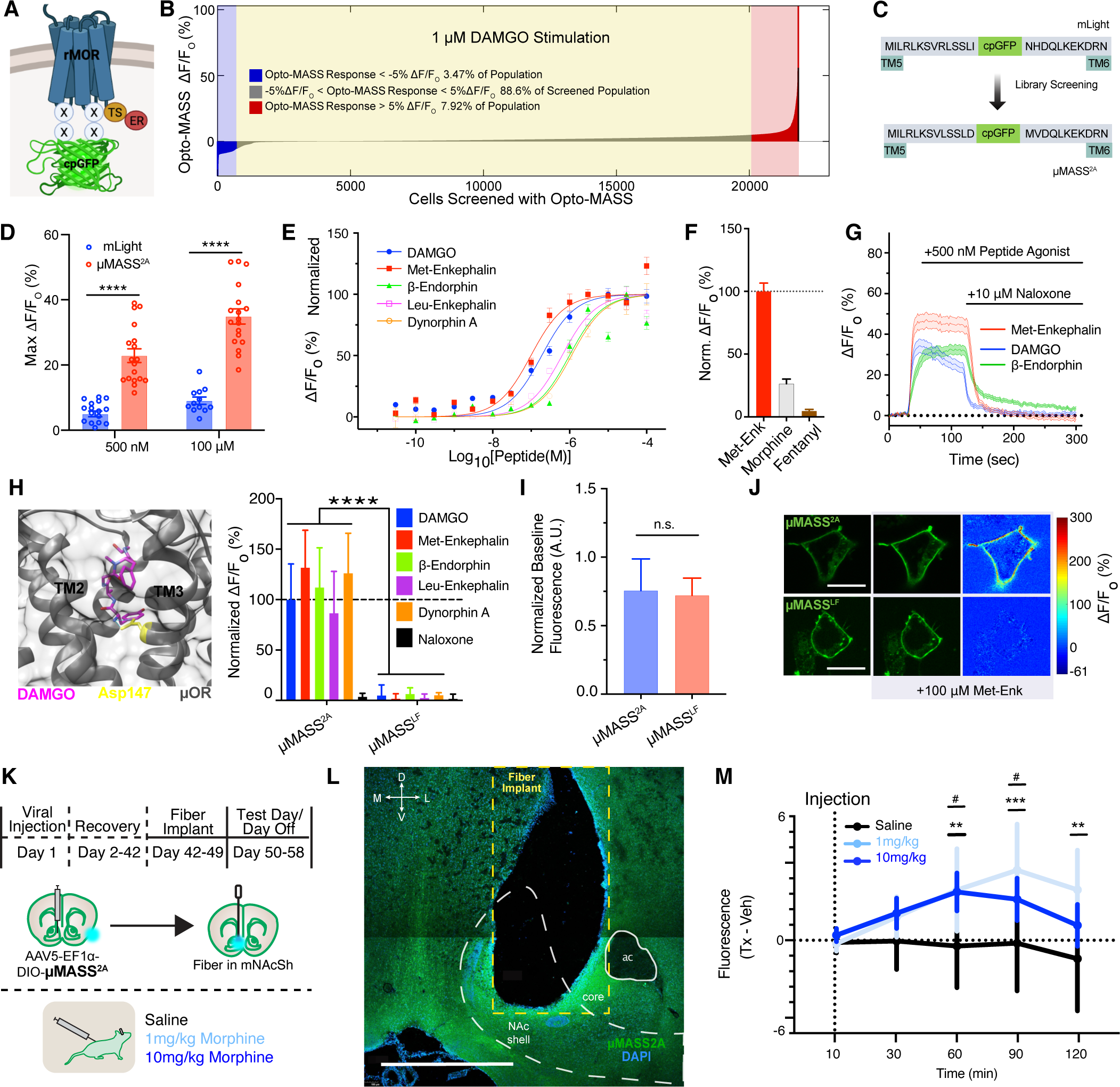
Design, Opto-MASS Optimization, and *in vitro* characterization of a neuropeptide sensor. A) cpGFP receptor insertion into rat mu-opioid receptor (MOR) and locations of linker residues targeted by site saturated mutation for generating a MOR sensor library. We added the membrane trafficking signals TS-ER to the c-terminus. B) Agregated sensor library response using Opto-mass. 21,839 cells were screened over two days, resulting in 3.47% with a ligand dependent ΔF/F_o_ < -5%, 88.6% with a ligand dependent -5% < ΔF/F_o_ < 5%, and 7.92% with a ΔF/F_o_ > 5%. C) Mutational changes in the selected variant μMass^2A^ compared to the parent scaffold mLight. D) mLight and μMASS^2A^ response to 500 nM (mLight 5.00±0.87%; μMASS^2A^: 22.9±2.1%) and 100 μM DAMGO (mLight 9.06±1.11, μMASS^2A^ 34.31±3.11). P <0.0001, unpaired t-test n = 12-18 cells, 2-3 wells. E) Apparent affinity curves of μMass^2A^ for common opioid peptides and DAMGO. K_D_ as follows: DAMGO 243 nM, Methionine-enkephalin 99 nM, leucine-enkephalin 637 nM, β-endorphin 1176 nM, Dynorphin A 1125 nM. n = 18 cells/3wells each. F) μMASS^2A^’s normalized response to 1 μM endogenous and exogenous opioids, morphine, and fentanyl. (n = 19-21 cells, three wells). G) μMASS^2A^ responses are reversible by addition of the opioid receptor antagonist naloxone. N = 27-28 cells/3 wells. H) Left: The mutated residue for generating a loss of function variant from μMASS^2A^ is highlighted in yellow (PDB ID: 6DDF). Right: Comparison of μMASS^2A^ and μMASS^LF^ (loss-of-function) ligand responses which were normalized to 10 μM DAMGO (Unpaired t-test, **** p<0.0001, n=3 wells, 6-9 cells per well). I) μMASS^2A^ and μMASS^LF^ do not have significantly different baseline fluorescence. The fluorescence was normalized to a C terminally tagged mRuby3 (p = 0.90 unpaired t-test with Welch’s correction, n = 16-27 cells/2-3 wells, mean ± SEM). J) Representative confocal images of μMASS^2A^ and μMASS^LF^ expressed in HEK293 cells and their responses to 100 μM met-enkephalin. K) Schematic of *in vivo* pharmacology μMASS^2A^ experiments. μMASS^2A^ was expressed in nucleus accumbens and fiber photometry recordings were obtained after injecting mice with saline, 1mg/kg morphine, or 10mg/kg morphine. L) Histological image of μMASS^2A^ expression and fiber tip placement in the nucleus accumbens. 1 mm scale bar. M) Mean fluorescence (circles) and SEM (bars) at 10, 30, 60, 90, and 120 minute time points after systemic injections of saline (black), 1mg/kg morphine (light blue) or 10mg/kg morphine (dark blue). * indicates statistical significance between 1mg morphine and saline doses. # indicates statistical significance between 10mg morphine and saline doses.

The exogenous opioid agonists morphine and fentanyl (at 1 μM) activated μMASS^2A^ at lower levels compared to Met-enk (26.4±3.64% and 4.63±1.23%, respectively) (Fig. 4F). We demonstrated that the sensor activity is reversible by applying the opioid receptor antagonists naloxone, which could abolish fluorescence signal and be used as an essential pharmacological control for *in vivo* experiments (Fig. 4G) [28]. Next, we engineered a loss of function (LF) variant by mutating Asp147^3.32^Gly. Asp147^3.32^ is hypothesized to coordinate a critical intermolecular bond with the primary amine in the canonical opioid signaling motif [29]–[31]. The μMASS^LF^ response signal to 10 μM of various opioid peptides was abolished (Fig. 4H). At the same time, it had a similar baseline fluorescence compared to the parent μMASS^2A^ (Fig 4I p = 0.90, unpaired t-test, Welch’s correction). Additionally, μMASS^LF^ lacks any membrane-localized fluorescence changes when imaged under confocal microscopy (Fig. 4J). Also, μMASS^2A^ did not have any significant ligand-dependent internalization in HEK293WT cells when exposed to 10 µM DAMGO and imaged over an hour at 37°C in comparison to a C-terminally tagged rat MOR (Supp. Fig 4E-G).

### *In vivo* opioid imaging with μMASS^2A^

Having developed an optimized GEFI for MOR, we next validated µMASS^2A^ *in vivo* in a brain site highly enriched in MORs, the nucleus accumbens (NAc). MORs in the NAc have long been shown to dramatically modulate motivated behaviors, including food consumption, social bonding, and drug seeking[19][32]–[34]. Although an important site for opioid reward, the temporal characteristics of opioid signaling *in vivo* have largely remained a mystery due to the inability to track it over subsecond timescales[24], [35]. Therefore, to determine whether µMASS^2A^ could monitor ongoing MOR activity, we used a combination of fiber photometry and pharmacological agonism (Fig. 4K, M). First, mice were habituated to a chamber where they could freely move around. During this period, µMASS^2A^ fluorescence was recorded to establish a relative baseline. µMASS^2A^ showed strong expression in nucleus accumbens (Fig. 4L). After 10 minutes, mice were injected with saline or 1mg or 10mg of morphine and allowed to continue exploring the chamber. The behavioral and fluorescent activity was recorded for a total duration of 2 hours. Overall, injections of 1mg and 10mg of morphine produced an increase in µMASS^2A^ fluorescence (Figure 4M) relative to saline test days (F_(8,32_= 1.7, P= 0.131). During the first 10 minutes of the test day, µMASS^2A^ activity was similar across drug conditions (t_1mg_= 0.998, p_1mg_= 0.06, t_10mg_= 0.904, p_10mg_= 0.40). The lower dose produced a more rapid response within 60 minutes of morphine injection (t_1mg_= 3.07, p_1mg_= 0.01) which persisted until the end of the two-hour period (t_1mg_= 3.78, p_1mg_= 0.001), although both doses were similar in magnitude by 60 minutes (t_1mg_= 3.07, p_1mg_= 0.01, t_10mg_= 2.96, p_10mg_= 0.01). Although 1mg and 10mg doses of morphine produce different maximal fluorescent responses, it should be noted that morphine only activates in µMASS^2A^ at≈25% of the capacity that met-enkephalin can produce. However, despite this ceiling, it should be appreciated that even the low dose of morphine was detectible by the biosensor, highlighting its sensitivity *in vivo*.

## DISCUSSION

We designed a novel high-throughput pipeline capable of engineering optogenetic sensors to detect ligands with distinctly different physiological roles *in vivo*. Opto-Mass provides several advantages over the field’s current methods. First, we harness a mammalian expression system that ensures only a single variant from our library is expressed in each cell. By expressing our libraries immediately in mammalian cells, we overcome problems presented by GPCR expression in *E. coli*. and yeast-based systems. Consequently, each screened variant passes the hurdle of mammalian transgene expression, protein folding dynamics, and trafficking. Additionally, using a doxycycline-inducible expression system ensures a controllable expression level for our constructs. We can use commercial transfection reagents to induce stable integration, removing the need for difficult viral packaging or dilution with ‘dummy’ plasmids with calcium phosphate transfection.

Second, due to time and resource restrictions, researchers usually functionally screen only hundreds or low thousands of variants for each sensor. In contrast, the Opto-MASS platform can functionally screen thousands of variants a day. While we did not screen our libraries’ entire possible mutational space yet, our throughput was still orders of magnitudes better than alternative methods. Thus, the unbiased approach is a step toward covering the mutational space of sensor libraries more efficiently and could help tp identify optimized sensor variants faster.

Third, the platform could easily be modulated to screen GEFI’s that are spectrally orthogonal to commonly used optogenetic actuators such as Channelrhodopsin2. For example, red-shifted sensors could be made with an eGFP control fluorophore. Optogenetic sensors can pair with actuators, like ChR2, for more precise experimental preparations. The development of optogenetic actuators, such as light-activated G_q_ proteins, could be developed by using calcium sensors in place of the control fluorophores.

Despite the advances Opto-MASS and the improved tools dMASS^3A^ and μMASS^2A^ present to the field, there is room for optimizations and further applications to the system. First, our gene recovery rate was around 30-40%, and further improvement would make the identification of high-performance genes easier. Second, we could expand our mutational libraries and functional screening space with sequential ligand addition. Third, while our high throughput screening of dopamine sensors did not shift the pharmacological profile of the sensor, screening with DAMGO may have biased the MOR-based sensor’s apparent affinity towards met-enkephalin over beta-endorphin. However, selecting different ligands for screening could pave the way for identifying variants with a different ligand specificity profile.

So far, we tested ∼26,000 dopamine and ∼23,000 mu-opioid sensor variants during pipeline optimization and library screening, which is less than 2% of the 20^4^ possible mutations in each case. However, this represents an order of magnitude higher throughput than previous studies. Importantly, this number was sufficient to identify already significantly improved variants. Furthermore, after transfection and expansion, we can freeze the library expressing cells in liquid N_2_ and recover them quickly before additional screening sessions (Fig.1). This effectively requires only one library generation, landing pad cell transfection, selection, and expansion step for each library. In contrast, traditional testing methods would require months if not years of preparations, measuring, and analysis and significantly more resources for DNA purification and cell cultures. We conclude that Opto-Mass’s enhanced throughput allows us to identify higher-performing constructs more efficiently by screening a significantly larger mutational space.

## Acknowledgements

A.B. was supported by The Brain Research Foundation, UW Royalty Research Fund, UW ISCRM IPA, NIGMS R01 GM139850-01, P30 DA048736-01-Pilot. K.A.M. was supported by NIGMS R35 GM142886, D.M.F. was supported by NHGRI RM1 HG010461, and NIGMS R01 GM109110. D.C.C. was supported by K99DA049862, and R00DA049862. G.D.S was supported by R01 DA032750, R01 DA038168, and P30 DA048736. The research received additional support from the Lynn and Mike Garvey Imaging Core, the UW NAPE Center, and ISCRM Shared Equipment. We want to thank Prof. Randy Moon for his support. Some figures were created with Biorender.com.

## Methods

### Microwell Array Fabrication

The master mold (negative) of the microwell arrays was fabricated using a ICP4-SPTS-DSi using deep reactive ion etching (DRIE) for etching silicon wafers. To prepare the wafer for DRIE, AZ1512 photoresist (MicroChemicals) was spun onto 100 mm silicon wafers (University Wafer) with a two-step process. First, wafers were spun at 500 RPM for 5 sec to spread the photoresist, with a final thirty-second spin step at 2500 RPM. After incubation at 100°C for 60 seconds on a hotplate, the wafer was exposed with a chrome on borosilicate glass mask and developed with AZ340 developer (MicroChemicals) [38].

Polydimethylsiloxane (PDMS) (Sylgard 184, Corning) was used to construct the microwell arrays. The elastomer and the curing agent were thoroughly mixed at a ratio 10:1 (w/w), poured onto the silicon wafer and placed in a desiccator to remove air bubbles. After the air bubbles were removed, the wafer was moved to a 55-75° C incubator to cure for several hours[38].

After curing, the PDMS was removed using razorblades and forceps. The PDMS was plasma treated for sixty seconds and then firmly pressed onto a dried bovine serum albumen (BSA) layer to nanostamp a layer of BSA onto the surface (FischerSci Cat# BP1600). The dried BSA layer was made by incubating a 2% w/v solution of BSA in PBS in a Petri dish for approximately 30 minutes. After, the dish was rinsed 3X with PBS and left to dry. After pressing the plasma-charged PDMS into the BSA, debris were removed from back of the PDMS using tape. The PDMS microwell arrays were then cut out using a scalpel and placed upright into a glass-bottom 24-well dish.

### Building Genetic Libraries for Screening on the Platform

Gibson Assembly was used to build the genetic libraries of sensors. The insert and backbone were PCR amplified using Q5 polymerase (New England Biolabs (NEB) M0515) and degenerate codons (IDT) were introduced with primers during PCR amplification of the cpGFP insert. To subclone the cpGFP moiety with two flanking mutational regions, we used primers to amplify the cpGFP domain out of a mammalian expression plasmid containing GCaMP6f. 1 µL of DpnI was used to digest PCR templates. PCR products were isolated from a 1% agarose gel stained with SyberSafe (Invitrogen Cat #S33102) and New England Biolabs (NEB) Monarch DNA Gel Extraction Kit (Cat # T1020L). After gel isolation, the insert and backbone were assembled using NEB HiFi DNA Assembly (NEB Cat #: E2621). A total of 0.2 pmol of DNA was used in the Gibson Assembly, with a 6:1 molar ratio of insert (cpGFP moiety) to vector and incubated at 50° C for 60 minutes.

The assembly was cleaned using a NEB PCR cleanup kit and double eluted from the column with 10 µL of prewarmed water. We then transformed 33 µl of electrocompetent cells (NEB Cat #C3020K) (2000 V, τ = 5ms) in ice-cold cuvettes (1 mm gap) with 2 µL of the elution. Immediately after pulsing the cells, 967 µL of prewarmed SOC media was added to the cuvettes. After 1 hour of recovery at 37° C and 240 RPM in a 15 mL recovery tube, a dilution of the recovery media was plated on an agar plate and grown overnight to estimate library size. The remaining recovery media was added to 125 mL of Luria Broth (LB) with ampicillin and grown overnight at 37° C and 240 RPM. The 15 mL recovery tube was rinsed several times with fresh LB media to ensure all transformants were added to the larger overnight culture.

After overnight growth, the agar plate was counted to estimate total transformants, and colonies were randomly selected for library sampling. Library plasmids were isolated using the Machery-Nagel NucleoBond Xtra Midi EF kit (Ref # 740420.50). The resulting plasmid prep was used to generate stably integrated cell lines with the landing pad cell line. The dopamine sensor library used dLight1.1 and the MOR sensor from reference [3]. During library screening, the MOR sensor template had an IgK sequence attached to the N terminus and the TSER signal to the C-terminus. The IgK sequence was removed from the μMASS^2A^ sensor during *in vitro* and *in vivo* characterization.

### Library Transfection into Landing Pad Cells

After validation, the library was correctly assembled in transformed *E*.*coli* through Sanger Sequencing selected colonies. HEK293T landing pad cells were stably recombined with our library using a double transfection protocol. Landing pad cells were maintained in standard growth media supplemented with 1-2 µg/mL doxycycline. On the day of transfection, the cells were gently lifted off the growth substrate using 0.05% Trypsin/EDTA (Invitrogen Cat # 25300120). Liftoff was stopped by adding growth media. Approximately 250,000 cells per well were seeded into 6-well dishes, and the final culture volume was 2 mL. The DNA transfection reagents were prepared by incubating 3 µg of plasmid DNA encoding the Bxb1 recombinase and 6 µL of Fugene6 reagent (Promega Cat # E2693) in 300 µL of Opti-MEM for 15 minutes and then added to the cell suspension. After 24 hours of incubation, cells were lifted off the growth substrate and centrifuged at 500 RCF for 5 minutes. After seeding at 250,000 cells/well in a 6-well dish, the cells were transfected a second time using the same protocol with library plasmids. For each round of library screening, five wells of the six well plate were transfected with the genetic library and combined after puromycin selection for screening.

### Cell Seeding

The nanostamped PDMS microwell arrays were then briefly plasma treated to charge the inner wells again. Quickly after plasma treatment, standard growth media was added to the wells, and they were placed in a desiccator to remove bubbles in the microwells. The plates were briefly returned to the tissue culture incubator to raise the temperature of the media and balance the pH. Next, the landing pad cells were lifted from the growth substrate with 0.05% Trypsin/EDTA. Once a single cell suspension was achieved, the cells were counted, and then 40K cells were seeded at a concentration of 0.5×10^6^ onto the arrays. Cells were slowly pipetted above the array with a micropipette. The cells were returned to the incubator for 10 minutes. After 10 minutes, the 24 well plates were then placed in a centrifuge and spun down at 100 RCF for 5 minutes. The arrays were then rinsed with growth media several times to remove cells not in microwells and cell debris. The final rinse is with DMEM/10% FCS supplemented with doxycycline at the concentration used during selection and half selection puromycin concentration. The cells were then returned to the incubator overnight.

### Library Screening and Cell Selection

On the morning of cell selection experiments, the arrays were washed twice with standard growth media and then once with imaging tyrode supplemented with GlutaMax (Gibco Ref: 35050-1), sodium pyruvate (GIBCO Ref: 11360-070), and MEM Non-Essential Amino Acids (Gibco Ref: 11140-050). A MetaMorph Journal was used to control the microscope during the imaging sequence. In brief, during the image capture stage, the arrays were imaged for the control fluorophore and then were imaged at GFP excitation/emission continually for one minute. Ligands were added by automatic pump or hand to the bath. The control fluorophore was imaged, and a bright-field image. The images were then analyzed using MetaMorph.

ROIs with the greatest response were identified and added to the live field of view and the image stack of the stimulation. The average intensity projection of baseline and stimulation images is divided to define the greatest responding ROIs. The image numbers that make up the ‘baseline’ and ‘stimulation’ are dependent on the time of ligand addition. The baseline images are typically defined as the first ten images of the stimulation time course. The proportion of images from the stimulation stack post-stimulation are selected to be a brief period after the ligand addition to the end of the stack.

The user can verify the selected ROIs prior to cell picking using glass micropipettes. The micropipette tips were then transferred to 200 µL PCR tubes with 5 µL of a TE/DTT buffer and immediately placed on dry ice for the remainder of the screening session. Positive pressure was applied during tip breakage into the solution.

After the screening session was over, the samples were processed to covert the mRNA of the sensor into cDNA using an adapted protocol of the ThermoFischer SuperScript IV Reverse Transcriptase (SSIV RT) protocol. 2 µL of the cDNA product was then amplified in a 25 µL reaction using Q5 polymerase. After DNA cleanup, the PCR product was Sanger Sequenced to check for contamination and cloned into a pCMV backbone using Gibson Assembly to validate the gene’s performance. After the transformation of the Gibson Assembly into chemically competent cells, the clones were grown up in 5 mL Luria Broth cultures with (100 µg/mL) ampicillin, and plasmid DNA was isolated using the Machery Nagel Endotoxin Free Miniprep kit. The plasmids were transfected into HEK293 WT cultures in plastic 24-well dishes to validate the performance of the gene.

### Reverse Transcriptase Reaction

Single-cell recovery tubes were prepared by diluting 5 µL of 0.1 M DTT into 200 µL of TE buffer. 5 µL of the TE/DTT buffer was added to each PCR tube. After a single cell was deposited into the tube, the tubes were incubated on dry ice for the remainder of the library screening session. After library screening, the tubes were removed from the dry ice and placed on wet ice. Each tube was processed with reagents from the SuperScript IV First-Strand Synthesis kit (Invitrogen Cat# 18091050). To each tube, 0.5 µL of 0.1 M DTT and 0.5 µL of RNAse Inhibitor were added. The samples were then placed on dry ice for five minutes and then moved back to wet ice. Next, 0.5 µL of the following was added to each tube, DI H_2_0, a 10 mM dNTP mix, and 2 µM primer.

Next, the primers were annealed to the mRNA by incubating the samples at 95°C for 30 seconds, 4°C for 1 minute, 65°C for 5 minutes. The samples were then returned to wet ice. Next, 2 µL of SSIV RT 5X Master Mix was added to each sample. The samples were pipetted up and down thoroughly. Finally, 0.5 µL of the SSIV RT Enzyme was added to each tube, and the samples were pipetted up and down thoroughly. The reverse transcriptase reaction was carried out in the following manner the samples were incubated at 53°C for 10 minutes and then at 80°C for 10 minutes to inactivate the reverse transcriptase. Next, 0.5 µL of RNAseH was added to each tube, and the samples were incubated for 20 minutes at 37° C to remove any mRNA from the cDNA. Samples were stored at -20° C before PCR amplification of cDNA with Q5 (New England Biolabs) or SuperFiII (ThermoFischer).

### Transfection for *In Vitro* imaging assays

HEK293 cells were cultured on tissue culture-treated plastic at 37° C with a 5% CO2 atmosphere. One day prior to transfection, cells were lifted off the growth substrate with 0.05% Trypsin/EDTA. The cells were then seeded into 24 well tissue culture plates. Cells were grown to 70-80% and then transfected. During transfection, growth media was replaced with fresh 250 µL of media. The DNA transfection reagents were prepared using the standard protocol. In brief, per well of transfection, 25 µL Opti-MEM, 1 μg of DNA, and 1.5 µL of P3000 were mixed. After five minutes of equilibration, the Opti-MEM/DNA/P3000 mix was added to a tube containing 25 µL Opti-MEM. And 1.5 µL of Lipofectamine. The DNA/P3000/Lipofectamine mix was incubated for approximately 15 minutes at room temperature before adding it to the wells. After incubation for 3-4 hours, the transfection media was removed and fresh media was added. Reactions were scaled for different wells according to the manufacturer’s directions.

Cells were imaged with an sCMOS camera (Photometrics Prime95B) on an epifluorescent microscope (Leica DMI8) using a 20X objective (Leica HCX PL FLUOTAR L 20x/0.40 NA CORR) forty-eight hours after transfection. A Lumencor Light Engine LED, and Semrock Filters were used for fluorescence imaging.

Before imaging, cells were rinsed once with tyrode (125 mM NaCl, 2 mM KCl, 2 mM CaCl_2_, 2 mM Mg Cl_2_ 30 mM Glucose, and 25 mM HEPES). Cells were imaged in Tyrode’s solution that was supplemented with GlutaMax (Gibco Ref: 35050-1), sodium pyruvate (GIBCO Ref: 11360-070), and MEM Non-Essential Amino Acids (Gibco Ref: 11140-050). Bath additions of ligands were made by hand for validation experiments and the volume added was always equivalent to the pre-addition bath volume. Ligands were prepared in a tyrode solution.

For confocal images of fluorescence responses, HEK293 cells were plated onto poly-l-lysine (Cultrex 3438-100-01) coated glass bottom plates and imaged in Tyrode’s solution with a Nikon A1R microscope with a 40X oil objective (CFI Plan Fluor NA 1.30) at room temperature (≈23°C). A 488 nm laser was used for GFP and sensor imaging and a 561 nm laser was used for red fluorophore imaging. Ligands were hand pipetted into the bath.

Fluorescence change graphs were generated by taking the average intensity projection of five frames before ligand addition and five frames after ligand addition in FIJI (NIH). The resulting images were then divided in MATLAB (2019a) to determine pixel-by-pixel fluorescence change, and a color scale was overlaid.

### Beta Arrestin Internalization Assay

12 mm glass coverslips were coated with poly-l-lysine. After incubation in poly-l-lysine for approximately one to two hours at room temperature or overnight at 4°C. After incubation, the coverslips were rinsed three times with 1X PBS. HEK293WT cells were seeded onto the coverslips and grown to 70-80% confluency. The HEK293WT cells were transfected using the above Lipofectamine 3000 reagents and protocol, with an increased total amount of DNA (1500 ng per well and a 1:1 molar split between the control fluorophore plasmids and sensor expression plasmids. The first contained the pCMV mRuby-CaaX, and the other contained a pCMV rMOR eGFP or the pCMV μMASS^2A^ plasmid.

Forty-eight hours after transfection, we imaged the coverslips on an epifluorescence microscope with a 40X oil objective at 37°C. Cells were imaged for one hour, during which they were sampled 60 times. A mCherry image and an eGFP image were taken at each time point. Images were analyzed in FIJI to collect intensity profiles, analyzed in Excel, and plotted in GraphPad Prism 8.

### Affinity Curves

HEK293WT cells were transfected with Lipofectamine 3000 reagents for the dopamine affinity curves. In brief, HEK293WT cells were seeded onto poly-L-lysine coated 96-well glass-bottom plates and expanded to 70-80% confluency and transfected. After transfection, the cells were incubated for 24-48 hours before imaging.

On the imaging day, cells were rinsed and imaged in 50 μL of supplemented tyrode. During imaging, 150 μL of a dopamine and tyrode solution was hand added to the bath during imaging. Cells were imaged with a 63X air objective.

For peptide affinity curves, landing pad cells expressing the μMASS^2A^ sensor were used. Cells were seeded at 25,000 cells per well in a 96 well dish and imaged the next day after overnight growth in 8 μg/mL doxycycline. A 63X air objective was used to image the cells continuously for sixty seconds. 150 μL of peptide solution was hand pipetted into a bath of 50 μL bath.

Cells were analyzed in FIJI (NIH). Five to six cells from each well were hand circled from background-subtracted image stacks (Rolling ball, 100 pixels). The average fluorescence intensity was measured for each cell and exported to Excel for analysis. Analyzed data were imported into GraphPad Prism 8 to calculate EC50 values using the nonlinear fit function and Least Squares fit.

### Calculation of ΔF/F_o_

The change in fluorescence was measured by hand circling regions of interest (ROIs) around background-subtracted images in FIJI. The ROIs were measured for mean grey value over time. The measurements were imported into Excel, where Equation 1 was used to calculate the change in fluorescence:

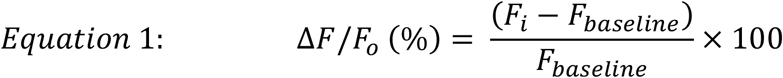

Where F_i_ is the ROIs mean fluorescence for a frame and F_baseline_ is the average fluorescence for the first five or six frames after imaging began. When peak fluorescence is shown, the maximum fluorescence is identified in a time course, and five frames surrounding maximum fluorescence are averaged to account for noise in the fluorescence measurement.

### Molecular Cloning

Unless explicitly stated, DNA constructs were cloned with either Q5 Polymerase, Platinum SuperFi II, Site-Directed Mutagenesis, Gibson Assembly, *In Vitro Assembly*, or standard restriction enzyme cloning. PCR products were verified on 1% agarose gels stained with SyberSafe and cleaned up with NEB Monarch PCR Clean-Up kits. The pCAG NLS HA bxb1 plasmid was gifted from the Fowler lab (Addgene #51271), the AttB puromycin plasmid used to generate libraries was gifted from the Fowler lab, and the pCMV dLight1.1 plasmid was sourced from Addgene (#111053), and rat MOR plasmid was a gift from the Bruchas Lab.

### *In Vivo* fiber photometry dMASS^3A^ and μMASS^2A^

Under isoflurane anesthesia (5-2%), 6 heterozygous Vgat-cre mice (9 weeks old, 3m and 3f) were injected with 400nL (Nanoject III, 1nL / minute) of AAV5-DIO-dMASS^3A^ into the dorsolateral striatum (AP: 1.25; ML: +/-1.6; DV: -2.3; angle: 10) and the contralateral nucleus accumbens (AP: 1.7; ML: +/-1.5; DV: -4.5 angle: 10) with hemispheres balanced across subjects. We then lowered 6mm optic fibers (Doric: 400µm, 0.37 NA, 1.25 zirconia ferrule) 0.1mm above each injection target and fixed the fibers to a head ring and the mouse’s skull using super glue and dental cement.

Following 1 week of recovery and return to presurgical body weight, mice were food restricted to 90% of their baseline body weight for 5 days before behavioral sessions. Mice were habituated to handling the head-fixation apparatus for 2 days before being head-fixed. The head-fixed behavioral apparatus consisted of a custom, 3d printed head fixation stage and 5x multi-spout that could rotate and retract using micro servos (Tower Pro SG92R). Each multi-spout was attached to an independent solenoid (Parker) which was calibrated before the experiment to ensure they delivered ∼1.5 µL per delivery. Licks were detected on each spout using a capacitive touch sensor (Adafruit MPR121). An Arduino Mega was used to control hardware and record the timing of behavioral and hardware events.

Mice were first trained to consume sucrose from a metal lickspout in a single 10-minute free-access session with a single spout in the extended position so that mice could freely lick for 30% sucrose. Free consumption was achieved by delivering a drop of sucrose via solenoid opening (∼1.5 µL over ∼15ms / delivery) immediately following each lick. Next, mice were trained on the forced-choice free-access multi-spout assay. Each session consisted of 100 trials of 3s access to 1 of 5 concentrations of sucrose (0, 5, 10, 20, 30%) with a random inter-trial interval of 11-16s drawn from a uniform distribution. During each access period, the spout extended forward, and the mouse could lick for sucrose for 3s, then the spout was retracted, and the multi-spout head was immediately rotated to the spout of the next trial. Mice were trained over 11 sessions and were then recorded over 3 sessions.

For μMASS^2A^ experiments, Penk-cre mice (9 weeks old, 2m and 3f) were injected with 200nL (Hamilton, 100nL/minute) of AAV5-EF1a-DIO-μMASS^2A^ into nucleus accumbens (AP: 1.7; ML: +/-1.0; DV: -4.4) and implanted with a 5mm optic fiber (Doric: 400µm, 0.48 NA, 5mm brass ferrule) 0.1mm above each injection target. Fibers were fixed to the skull with Metabond.

Following 6 weeks of recovery, mice were habituated to the test chamber (25cm x 25cm x 25cm) and allowed to explore for 30 minutes. Near the end of the habituation day, mice were systemically injected with saline (i.p.) to habituate them to the injection procedure. 48 hours later, mice were again placed into the test chamber and allowed to explore for 10 minutes. After 10 minutes, mice were injected with saline or 1mg/kg or 10mg/kg morphine and returned to the test chamber for the remaining duration of the two-hour test. Behavioral videos and photometry recordings were collected for the entire two-hour test.

At the conclusion of the experiment, mice were transcranial perfused with 20mL of PBS and 20 mL of 4% PFA. Skulls were removed and postfixed for 24h before the brain was removed and postfixed for an additional 24h. Brains were frozen at -20°C and then sectioned at 40µm on a cryostat (Leica). Sections were collected in PBS, then mounted onto glass slides, and coverslips using fluoroshield with DAPI (Sigma) for staining. Sections were imaged at 5x magnification under an epifluorescence microscope (Zeiss ApoTome2) using Zen (Blue Edition, Zeiss) or at 10x magnification under a confocal microscope (Olympus Fluoview FV3000) using FV31S-SW. The location of optic fibers was determined by mapping fiber position onto a mouse histological atlas (The Mouse Brain, Paxinos 2001).

We recorded dMASS^3A^ fluorescence in the NAc and DMS simultaneously by connecting each fiber to patch cables (Doric: 400µm, 0.37 NA, 1.25 zirconia ferrule) coupled to a 5-port mini cube (Doric) and integrated fiber photometry system (RZ10X, Tucker-Davis Technologies). We used 465nm light modulated at 331 Hz for measuring dMASS^3A^ fluorescence and 405nm light modulated at 209 Hz for measuring autofluorescence. Light emission was collected using the same fiber and was measured using a photosensor (Lux). During collection, signals were low pass filtered at 6Hz and demodulated. Excitation power for both wavelengths was set to 30 µw. The timing of hardware and behavioral events were recorded using TTL inputs to the fiber photometry system.

For μMASS^2A^ experiments, we recorded μMASS^2A^ fluorescence in the NAc by connecting an optic fiber to the implanted fiber using a ferrule sleeve (Doric, catalog no. ZR_2.5). Two light-emitting diodes (LEDs) were used to excite μMASS^2A^. A 531-Hz sinusoidal LED light (Thorlabs, LED light, catalog no. M470F3; LED driver, catalog no. DC4104) was bandpass filtered (470 ± 20 nm, Doric, catalog no. FMC4) to excite μMASS^2A^ and evoke μ opioid-dependent emission. Laser intensity for the 470-nm wavelength band was measured at the tip of the optic fiber and adjusted to 50 μW before each day of recording. μMASS^2A^ fluorescence traveled through the same optic fiber before being bandpass filtered (525 ± 25 nm, Doric, catalog no. FMC4), transduced by a femtowatt silicon photoreceiver (Newport, catalog no. 2151), and recorded by a real-time processor (TDT, catalog no. RZ5P). The timing of the injection was recorded using behavioral video recordings. The envelopes of the 531-Hz signal were extracted in real-time by the TDT program Synapse at a sampling rate of 1,017.25 Hz.

Fiber photometry signals for dMASS^3A^ were post-processed using custom Python and R scripts. The 405nm channel was inspected for abrupt changes in signal power that could be attributed to motion, but none were observed (probably because the mice were head-fixed throughout the recording). As a result, the 405nm channel was not used to correct the dMASS^3A^ signal and was discarded from further analysis. We corrected for photobleaching for each session by fitting and subtracting a 4^th^ degree polynomial. We normalized the fluorescent signal by taking a z-score using the mean and standard deviation of the signal throughout the entire session. The signal was then smoothed using a 100ms moving average and then downsampled to 20Hz. Next, we used behavioral time stamps to extract peri-event time histograms centered on the access period and then baseline corrected by subtracting the mean signal during the 3s before the onset of access.

For μMASS^2A^ experiments, fiber photometry signals were post-processed using custom MATLAB scripts. We corrected for photobleaching for each session by fitting our 470 fluorescent signals to a 4-term polynomial function. We normalized the fluorescent signal by subtracting the vehicle-treated fluorescent signal from the treatment group signal. Next, we used behavioral video recordings to extract the time of injection and baseline corrected by subtracting the mean signal during the 9.5 minutes before injection. To calculate the fluorescence at the desired time points, we averaged a range of the raw, baseline corrected, decay adjusted 470 fluorescence from +/- 1 minute from the desired timepoint (ex. average fluorescence from 9 to 11 minutes for 10-minute time point) for each animal and treatment group.

## Notes

### Competing Interest Statement

The authors have declared no competing interest.

### Summary of Updates

1. We added Carrie Stine as a co-author 2. Fig. 2K, right panel: We changed the y-axix label from "Opto-MASS dF/F0 (%)" to "Opto-MASS CoV (%)"

