## Supplemental Figures for "Opto-MASS: a high-throughput engineering platform for genetically encoded fluorescent sensors enabling all-optical *in vivo* detection of monoamines and opioids"

#### Supplemental Figure 1

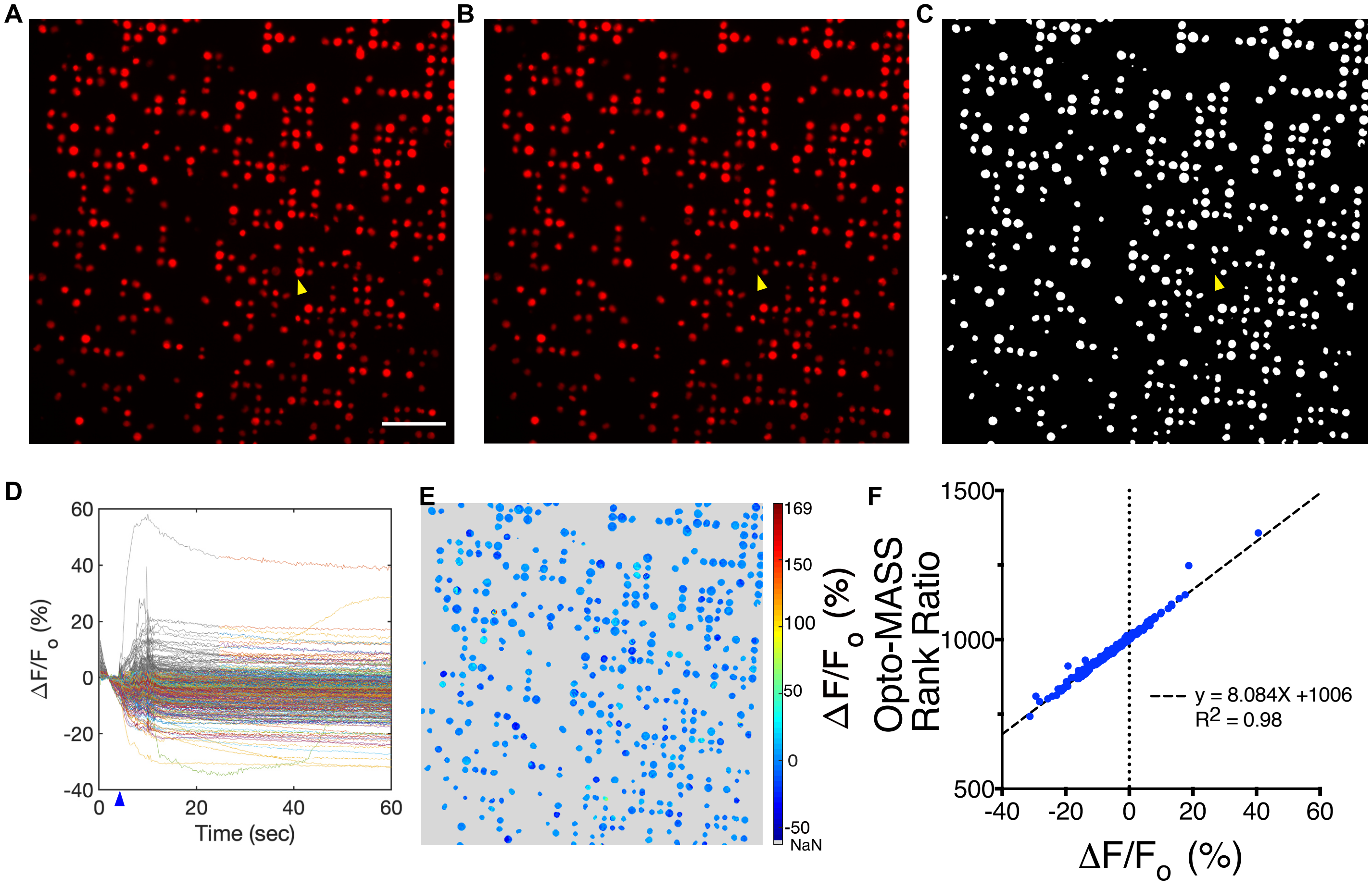

#### Supplemental Figure 2

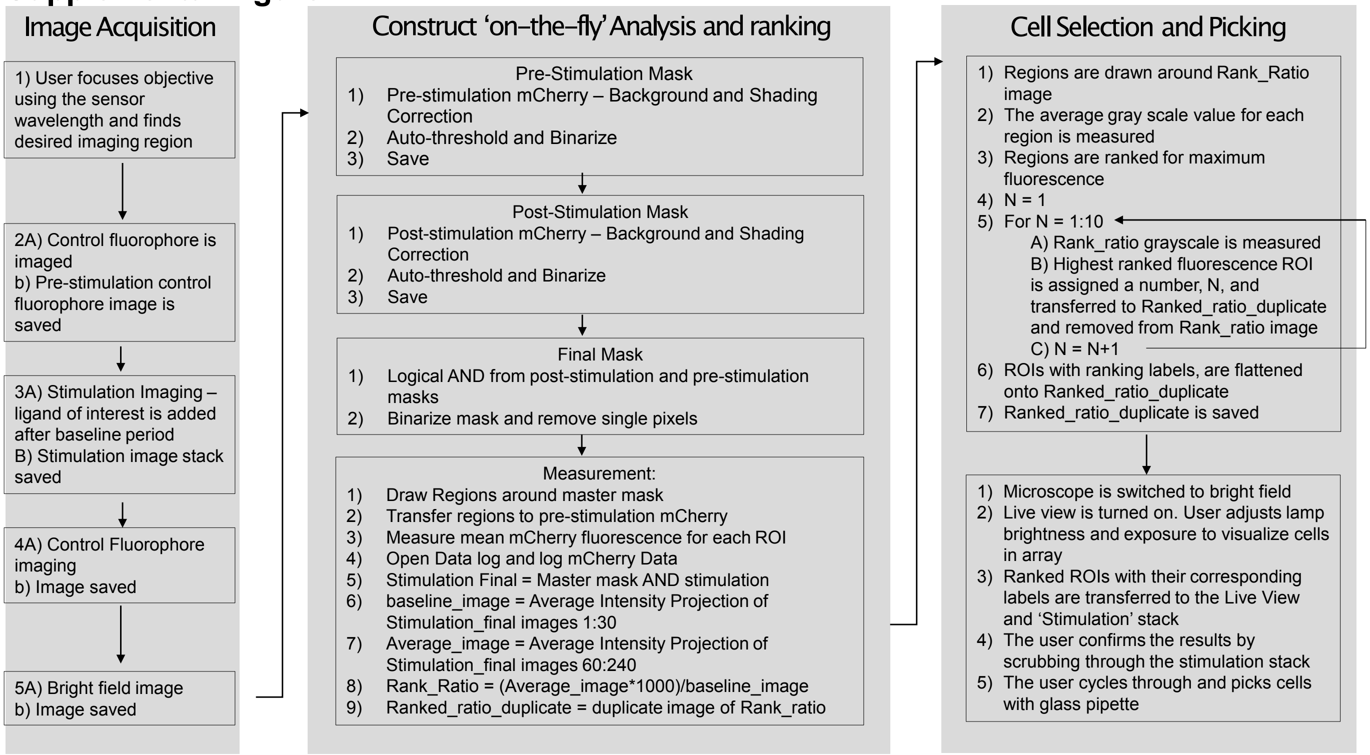

#### Supplemental Figure 3

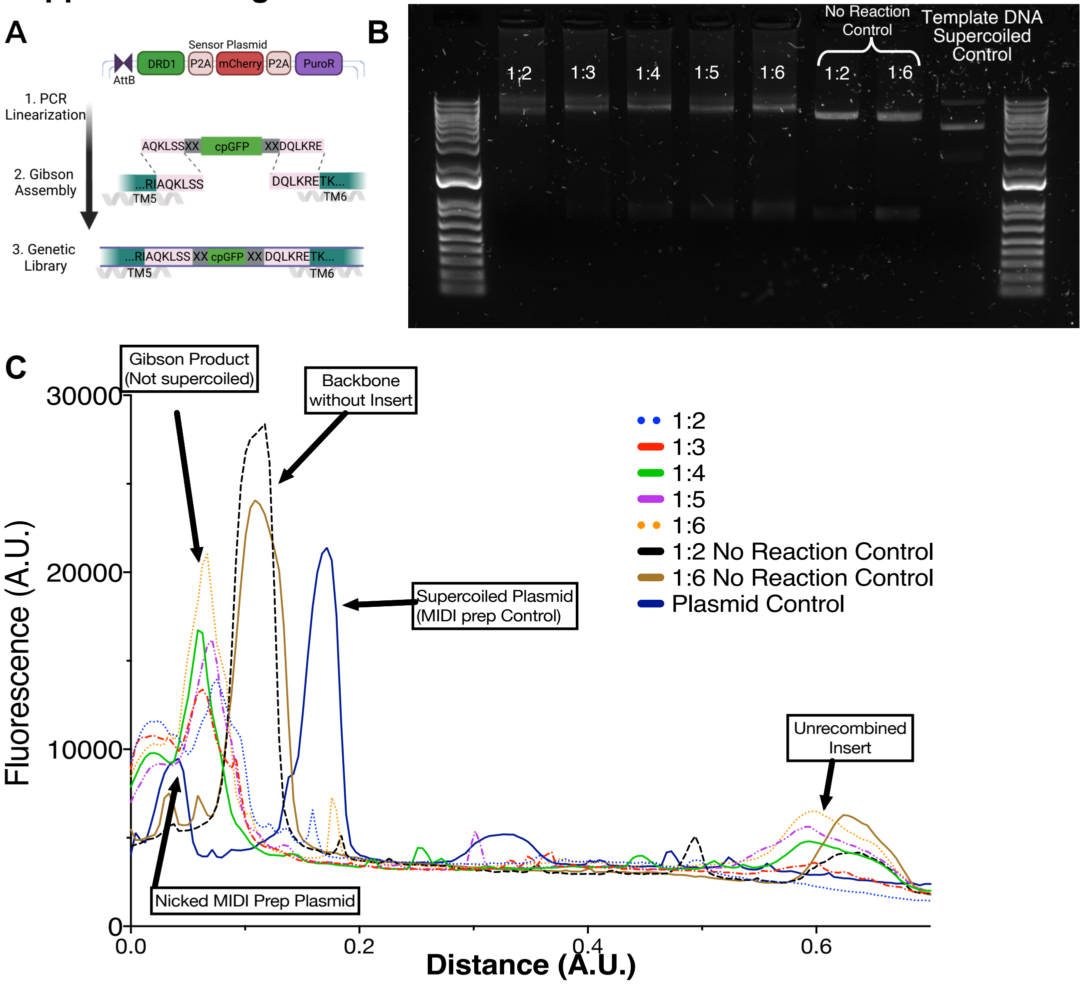

### Supplemental Figure 4

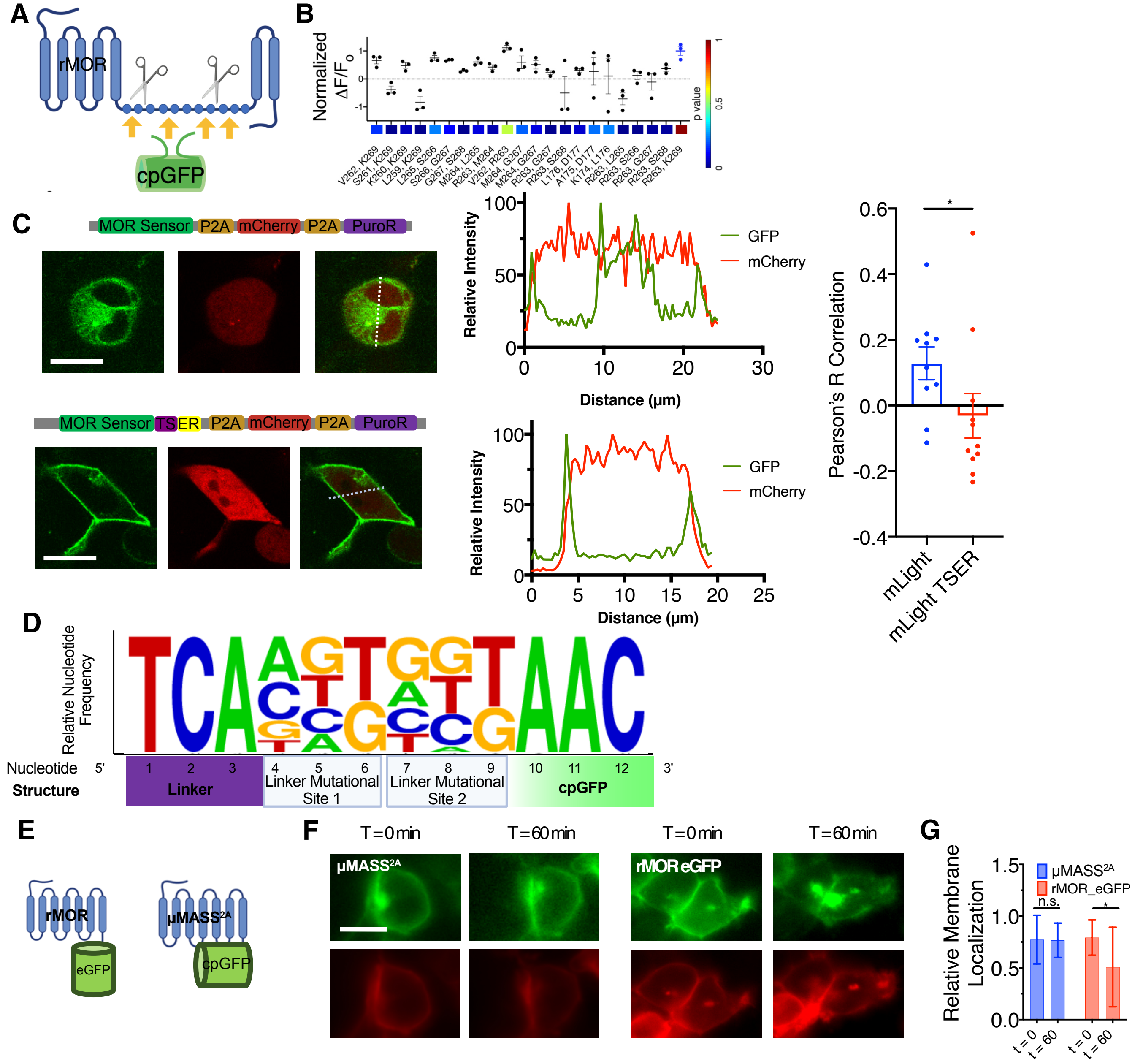

**Supplemental Figure 1: A rapid analysis algorithm to identify high-performance sensors**

A) Example of a false-color image of the negative control fluorophore mCherry pre-stimulation. Yellow arrow denotes an ROI that moved during stimulation. Scale bar 200 μm.

B) Post-stimulation imaging of mCherry of the same array. Yellow denotes the absence of the ROI highlighted in A.

C) Binary mask generated from pre and post-stimulation images in A and B. Note the lack of an ROI for the excluded cell, denoted by the yellow arrow.

D) Individual time-resolved traces of green fluorescence dynamic extracted from the generated ROI mask. Cells were stimulated by 500 nM dopamine. The Blue arrow denotes the approximate time of DA addition. Signals during the presence of motion artifacts due to DA addition were excluded.

E) Heat map of pixel intensity changes in the ROI mask upon ligand addition. Note that areas outside the mask were assigned a grey value.

F) The Opto-MASS rank ratio generated by the division of the baseline and post-stimulation average intensity projections correlates with the post-hoc calculation of fluorescence change. Note that only the average fluorescence change for the last 35 seconds was used to measure a change in fluorescence percentage. This excludes earlier motion artifacts.

**Supplemental Figure 2:** Detailed overview of high throughput screening execution.

**Supplemental Figure 3: Optimization of Library Generation**

A) Schematic of the dopamine sensor library construction using Gibson Assembly. (HiFi DNA Assembly Master Mix, NEB # E5520).

B) 1% agarose TAE gel. Lanes, from left to right: backbone:insert ratios: 1:2, 1:3, 1:4, 1:5, 1:6; 1:2 and 1:6 no reaction control and supercoiled DNA of the parent scaffold used for the library. The assembled plasmid travels slower than the linear backbone through the agarose due to not being supercoiled (see lane 1:2 Gibson Product). The molar amount of backbone added to each Gibson Assembly reaction was held constant. 1:6 molar ratio was used for library construction.

C) Fluorescence analysis of the lanes, averaged over 30 pixels wide. Image analysis was done in FIJI. Correctly assembled Gibson products will travel slower than super coiled plasmid DNA of the template.

**Supplemental Figure 4: Design, Engineering, and Characterization of μMASS Libraries and extended biophysical characterization of improved variants**

A) Schematic showing the insertion of the cpGFP reporter domain into the rat mu-opioid receptor.

B) cpGFP domain insertion screening results. Residues denote the position of the N- and C-terminal MOR residues next to the cpGFP linker. The original insertion of cpGFP in mLight between R263 and K269 yielded the highest signal amplitude. Three wells, ANOVA, multiple comparisons.

C) Membrane trafficking signals (TS-ER) were added to the MOR sensor to reduce cytotoxicity from aggregation in the cytosol in the HEK293T landing pad cell line. Cells were imaged on a 40X confocal microscope and analyzed for low correlation with the cytosolic mCherry fluorophore. Summary data n = 2 wells, 5 cells/well. Mann-Whitney Test, p =0.0357.

D) Relative nucleotide frequency in eighteen selected colonies from the μMASS library generation. Logo creation using [36], [37].

E) rMOR-EGFP and $\mu MASS^{2A}$ internalization experimental set up.

F) Representative epifluorescent images of HEK293 cells expressing membrane-localized control fluorophores (red, mRuby-CAAX) and either rMOR tagged with eGFP or green fluorescent $\mu MASS^{2A}$. Images were taken before adding 10 uM DAMGO and 60 min after. 10 um scale bar.

G) Summary data for the relative membrane localization of GFP fluorescence before and after one hour of 10 uM DAMGO addition. rMOR internalizes while $\mu MASS^{2A}$ remains localized in the plasma membrane. N = 2-3 wells, 4-5 cells/well. Paired t-test, n.s. p>0.05, p=0.035 rMOR_eGFP.
